# Genome-wide chromatin reset at the onset of leg regeneration in pancrustaceans

**DOI:** 10.64898/2026.10.09.757970

**Authors:** Maëlle Louvier, Gillian Forbes, Mahmood Mohammed-Ali, Eve Moutaux, Florent Chuffart, Arthur Argoud, Fabien Girard, Agnès Dumont, Belén Pachano, Christopher Swale, Mohamed-Ali Hakimi, Romain Koszul, Aurèle Piazza, Michalis Averof, Mathilde Paris, Guillermo A. Orsi

**Affiliations:** Institute for Advanced Biosciences - IAB, Université Grenoble-Alpes, INSERM U1209, CNRS UMR 5309, Grenoble, France; Institut de Génomique Fonctionnelle de Lyon (IGFL), École Normale Supérieure de Lyon, CNRS UMR 5242, Université Lyon 1, Lyon, France; Institut Pasteur, CNRS UMR 3525, Université Paris Cité, Unité Régulation Spatiale des Génomes, Paris, France; Laboratoire de Biologie et Modélisation de la Cellule (LBMC), École Normale Supérieure de Lyon, CNRS UMR 5239, Université Lyon 1, Lyon, France

**Author notes:** equal contributions (Gillian Forbes, Mahmood Mohammed-Ali).

## Abstract

Regeneration requires injured tissues to selectively reactivate certain developmental programs while triggering regeneration-specific mechanisms and responding to the immediate consequences of tissue damage. How chromatin states are reorganized during this transition, and whether the underlying sequence of regulatory events is conserved across regenerating animals, remain poorly understood. We address these questions in the two-spotted cricket *Gryllus bimaculatus*, in which limb regeneration progressively declines during nymphal development and is absent in adults, and extend our analysis to the crustacean *Parhyale hawaiensis*. We first establish a chromosome-scale genomic and epigenomic framework for the cricket and then follow chromatin and transcriptional changes after limb amputation. Amputation triggers a striking genome-wide shutdown of enhancer-associated chromatin, marked by loss of accessible regulatory regions and depletion of H3K27ac and H3K4me1. This is followed, in regeneration-competent tissue, by selective reopening of a restricted population of regulatory elements and strong induction of the pioneer transcription factor Zelda (GbZelda). The GbZelda locus is unusually marked by both H3K27me3 and H3K9me3 in intact leg tissue; after injury, H3K9me3 is reduced broadly, whereas H3K27me3 is selectively relieved in regenerating nymphs. Finally, analysis of *Parhyale* reveals the same sequence of chromatin events during appendage regeneration: an initial enhancer shutdown followed by selective regulatory reopening concomitant with Zelda upregulation. Thus, rather than restoring the pre-injury regulatory landscape wholesale, appendage regeneration in these two pancrustacean models proceeds through a conserved sequence of genome-wide chromatin resetting, selective regulatory reopening and activation of a pioneer transcription factor.

## INTRODUCTION

The capacity to regenerate lost or damaged body parts varies widely among animals, and regenerative competence is frequently restricted during development and lost in adulthood (Tanaka et al., 2003; Seifert et al., 2023; Alibardi, 2024; Yun, 2015; Cai et al., 2023). Understanding how an injured tissue re-enters a developmentally plastic state capable of rebuilding a complex structure is therefore a central question in regenerative biology. In particular, it remains unclear how the chromatin landscape is reorganized after injury, how developmental regulatory programs are re-engaged, and whether regeneration involves similar chromatin remodelling processes in distantly related animals.

Arthropods provide particularly useful systems for addressing these questions because appendage regeneration is closely integrated with developmental progression and molting (Das, 2015; Bando et al., 2018; Suzuki et al., 2019; Zhong et al., 2023). Crustaceans, insects and chelicerates can regenerate appendages to varying extents, and regeneration typically combines wound healing with blastema formation, patterning and subsequent growth. Among insects, the two-spotted cricket *Gryllus bimaculatus* is well suited for studying this problem because it regenerates amputated legs during nymphal development but not after the adult molt (Mito and Noji, 2008; Nakamura et al., 2008; Bando et al., 2018). Previous studies have implicated Wnt, JAK/STAT, EGFR, Hippo and other pathways in cricket leg regeneration and have revealed the cellular dynamics accompanying blastema formation and regrowth (Niwa et al., 2000; Bando et al., 2013; Bando et al., 2022). Chromatin regulation is also involved: depletion of H3K27me3 through knockdown of the Polycomb methyltransferase GbE(z) perturbs blastema development and limb regrowth (Hamada et al., 2015).

The chromatin changes associated with regeneration have been studied in a number of organisms. Injury can reactivate developmental enhancers, regeneration-specific enhancers and embryonic transcription factors, while other damage responses involve rapid activation of regulatory elements controlled by factors such as AP-1 (Kang et al., 2016; Wang et al., 2020; Harris et al., 2020; Harris, 2022; Kawaguchi et al., 2024). These observations have led to the view that, beyond activating regenerative programs, regeneration may require cells to escape the chromatin states that stabilize differentiated identity. Indeed, chromatin constitutes a barrier to cellular reprogramming, both in vitro, where disruption of CAF-1 or heterochromatin facilitates acquisition of alternative cell states (Soufi et al., 2012; Chen et al., 2013; Cheloufi et al., 2015), and in vivo, where partial reprogramming can enhance cellular plasticity and regenerative capacity (Chen et al., 2022), and where remodeling of H3K36 methylation promotes plasticity and regeneration following intestinal injury (Pashos et al., 2025). Thus, regeneration may involve not only activation of regenerative elements, but also a transient resetting of chromatin constraints. Yet it remains unclear whether injury first induces such a broad reorganization of chromatin, subsequently resolved into a regeneration-specific state, or instead selectively activates regeneration-associated elements.

These questions also raise an evolutionary question. The capacity for appendage regeneration is widespread across crustaceans and insects (Pancrustacea), yet it is not known whether similar chromatin transitions are associated with regeneration in different species. An informative comparison with crickets is provided by the crustacean *Parhyale hawaiensis*, in which appendage regeneration can also be studied at genomic scale (Sinigaglia et al., 2022; Paris et al., 2022).

In this study, we address how chromatin is reorganized during appendage regeneration and whether the regulatory events accompanying regeneration are shared across pancrustaceans. First, we combine transcriptomic and chromatin profiling of nymphal (regenerating) and adult (non-regenerating) cricket legs to characterize the dynamics of gene expression/regulation following amputation. We find that injury is initially accompanied by a genome-wide shutdown of enhancer-associated chromatin, characterized by loss of chromatin accessibility and the active enhancer-associated histone modifications H3K27ac/H3K4me1, followed by selective reopening of a restricted set of regulatory elements in regenerating tissue. This later transition coincides with strong upregulation of the pioneer transcription factor Zelda (GbZelda), whose locus is controlled by an unusual combination of H3K27me3 and H3K9me3, with selective release of both marks in regenerating nymphs. Finally, analysis of regenerating *Parhyale* legs provides evidence of a similar sequence of enhancer shutdown, selective reopening and Zelda upregulation during the course of leg regeneration. Together, our results identify a conserved sequence of chromatin reorganization during pancrustacean appendage regeneration, in which a genome-wide loss of regulatory activity is followed by selective reconstruction of a Zelda-associated regulatory landscape.

## RESULTS

### Experimental framework for studying limb regeneration and chromatin dynamics in *G. bimaculatus*

*G. bimaculatus* undergoes eight nymphal stages before the adult molt (Mito and Noji, 2008; Nakamura et al., 2008; Bando et al., 2018). Injured legs can be regenerated during the nymphal stages, but this capacity is lost in adults. We sought to quantify this decline in regenerative capacity, and evaluate the extent to which growing nymphs are regeneration-competent. To establish a baseline for normal growth, we measured body and tibia size throughout nymphal development (Supplementary Figure 1). We then amputated the tibia of the third thoracic leg at approximately its midpoint in stage 3, stage 5 and stage 7 nymphs, and in adults (Fig. 1A,B). The growth of the amputated leg was then compared with that of the contralateral intact leg, through the subsequent molts, allowing regeneration efficiency to be normalized to the substantial growth that occurs independently of regeneration. These measurements reveal a progressive decline in regenerative efficiency (Fig. 1B). After two molts, regenerated legs reach approximately 78% of the size of contralateral legs in amputated stage 3 animals, 62% in stage 5 animals, and 51% in stage 7 animals. Amputated adult legs fail to regrow after a similar period of time. Thus, the loss of regenerative ability is not associated with an abrupt transition during the adult molt, but shows a progressive reduction during nymphal development.

**Figure 1.**
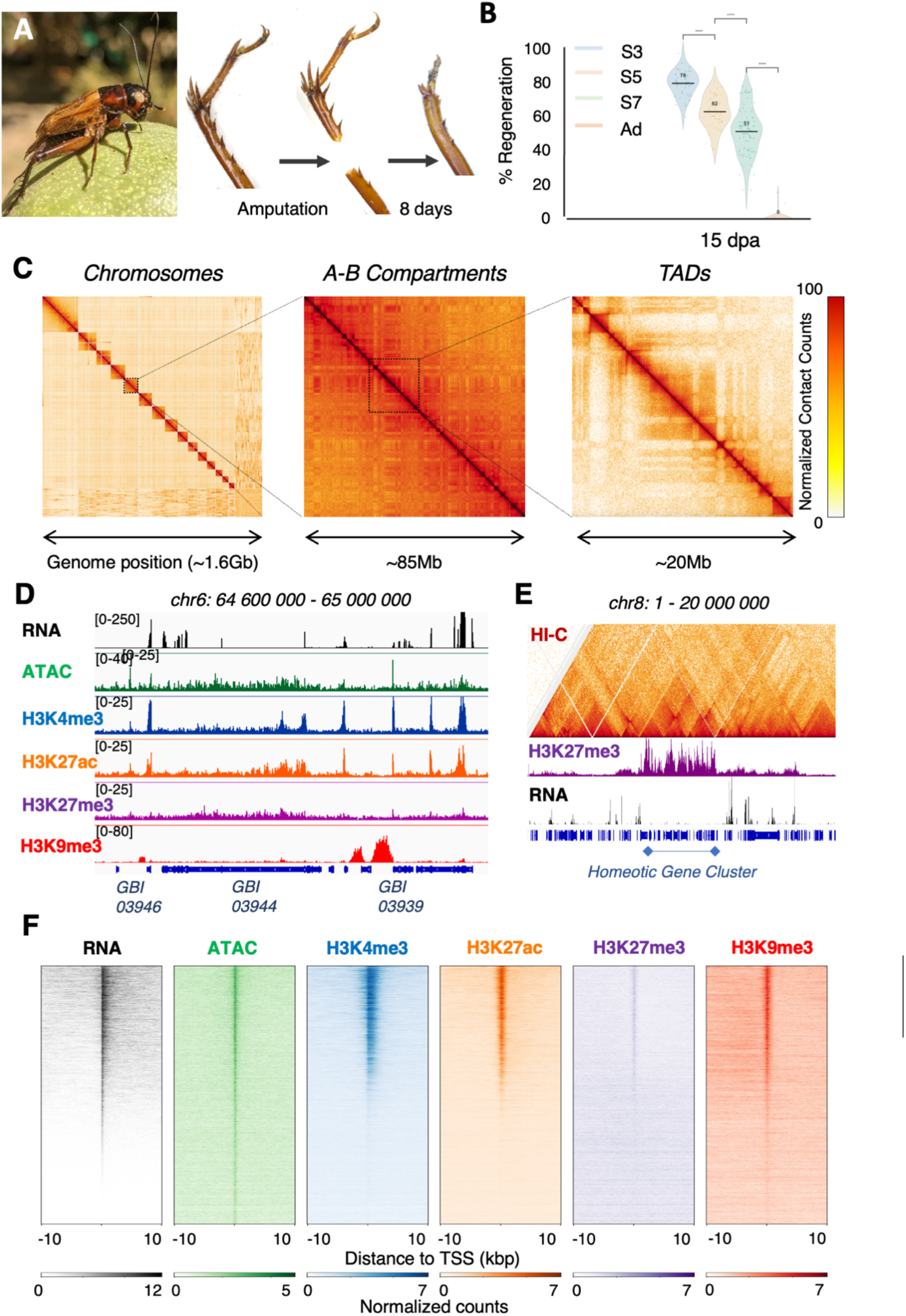
Developmental and epigenomic framework for studying leg regeneration in the cricket *Gryllus bimaculatus*. (A) Cricket limb regeneration after amputation. (B) Quantification of regeneration efficiency (see Methods) in nymphal stages 3, 5 and 7 and in adults. (C) Hi-C maps illustrating chromosome-scale organization, A/B compartments and TADs. (D) Example genomic region showing RNA, ATAC-seq and histone-modification profiles. (E) Example of a broad H3K27me3 domain containing the homeotic gene cluster. (F) Aggregate chromatin profiles around transcription start sites (see Methods).

Next, we sought to establish the genomic resources necessary for studying chromatin changes during regeneration. We generated a chromosome-scale assembly of the *G. bimaculatus* genome by combining long-read Nanopore sequencing, Hi-C and existing genome sequence data. The resulting assembly is approximately 1.6 Gb, with more than 87% of the sequence assigned to the 15 chromosomes, comprising 14 autosomes and the X chromosome (Fig 1C). Hi-C contact maps display preferential contacts within individual chromosomes, a checkerboard pattern characteristic of A/B compartmentalization and boundary-delimited topologically associated domains (TADs). These features are broadly consistent with the chromosome-level organization reported in other metazoans and in recently published *G. bimaculatus* assemblies (Bickmore and van Steensel, 2013; Dixon et al., 2012; Dekker et al., 2017; Kataoka et al., 2026; Li et al., 2026)(Supplementary Figure 2).

Having established the genome organization, we next characterized the distribution of chromatin accessibility and histone modifications in relation to transcription in the cricket genome. Ovaries provided a readily accessible source of abundant tissue for this initial assessment of our chromatin profiling approach (Fig 1D-F). To visualize chromatin patterns around gene promoters and regulatory elements, we aligned genes at their transcription start sites and plotted the average signal for each chromatin assay across the surrounding regions (Fig. 1F, Supplementary Fig. 3). These aggregate profiles represented as heatmaps highlight patterns shared across genes rather than the signal at individual loci. RNA-seq and ATAC-seq, together with H3K4me3 and H3K27ac profiling, revealed accessible chromatin and active transcription-associated histone marks at actively transcribed promoters and putative regulatory regions (Fig. 1D–F; Supplementary Fig. 3).

The H3K27me3 mark, associated with Polycomb repression, showed modest enrichment at many promoters as well as broad domains spanning multiple genes. One prominent domain encompassed the entire Hox locus, which was transcriptionally inactive in ovaries, consistent with its expected repression in this tissue (Fig. 1E). The constitutive heterochromatin H3K9me3 mark was also preferentially enriched at active promoters extending into upstream non-coding regions at a subset of active genes (Fig. 1D,F; Supplementary Fig. 4). Upstream regions with greater H3K9me3 enrichment showed lower transcription compared to upstream regions with less enrichment. This association raises the possibility that H3K9me3 restricts upstream transcription, although these data do not establish a causal role. Together, these experiments provide an initial characterization of cricket chromatin organization and support the application of our profiling approach to the response to leg amputation.

### Amputation induces a genome-wide enhancer shutdown independently of regenerative competence

Next, we interrogated chromatin dynamics shortly after leg injury in stage 7 nymphs, which retain substantial regenerative capacity, and in adults, which do not regenerate. Our goal was to distinguish chromatin changes that are associated with regeneration from changes that follow tissue damage in the absence of regeneration. In both stages we amputated the third thoracic leg, at the tibia, and collected approximately 1 mm of tissue from the tip of the remaining stump three days after amputation (Fig. 2A). For each condition, similarly sized and located sections of tissue from the contralateral intact legs provided stage-matched controls.

**Figure 2.**
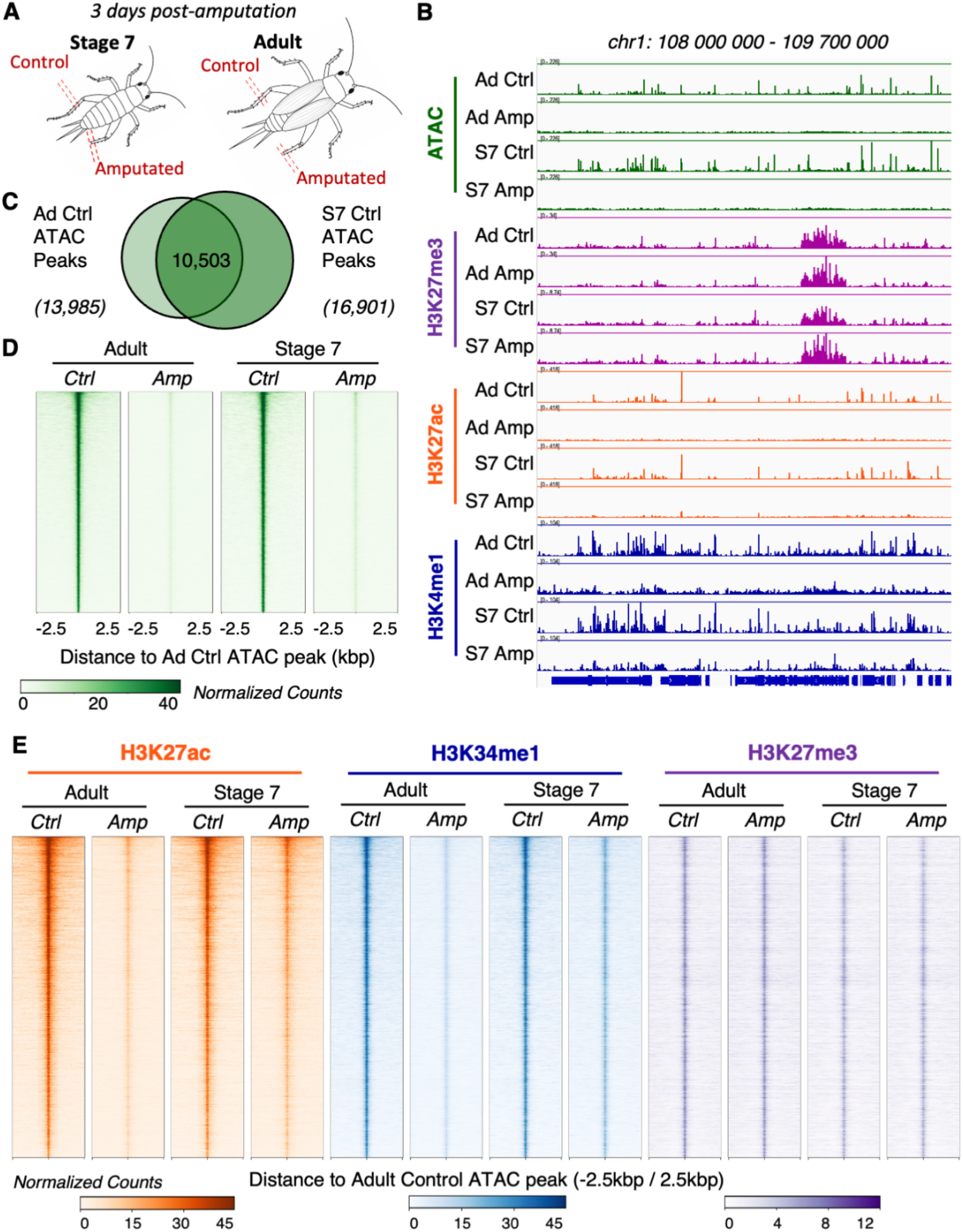
Limb amputation leads to a genome-wide enhancer shutdown independently of regenerative competence. (A) Experimental design comparing control and amputated stage 7 and adult legs at 3 days post-amputation. (B) Genome-browser profiles illustrating the loss of ATAC-seq signal and enhancer-associated histone marks after amputation. (C) Overlap between adult and stage 7 control ATAC-seq peaks. (D) Enrichment of ATACseq signals around ATACseq peaks identified in adult control samples. (E) Aggregate profiles of H3K27ac, H3K4me1 and H3K27me3 signals around ATACseq peaks identified in adult control samples.

We first examined chromatin accessibility using ATAC-seq. In control legs, the profiles of stage 7 and adult legs are very similar, with discrete accessible regions distributed throughout the genome, with substantial overlap in the profiles of nymph and adult legs (Fig 2B, C). In amputated leg stumps from both stage 7 nymphs and adults, the discrete ATAC-seq peaks that characterize the control legs were strongly reduced, and the chromatin accessibility profiles became much more homogeneous across the entire genome (Fig 2B, D). The effect was not restricted to a small group of loci: the overall landscape of accessible chromatin was profoundly altered across the genome. Only a small number of loci retained opening, and this was only partial, while we did not detect any amputation-specific peaks (Fig 2B and Supplementary Figure 5).

We then asked whether other chromatin features associated with enhancer activity showed the same behavior. CUT&Tag profiling for H3K27ac and H3K4me1, two canonical enhancer-associated histone modifications, revealed a similarly strong depletion of signal at regulatory regions after amputation (Fig. 2B, D). Importantly, this depletion was observed in both stage 7 and adult legs. In contrast, when we used CUT&Tag to profile the distribution of H3K27me3, which is normally associated with stable Polycomb-dependent repression, we found that this histone mark is comparatively stable between intact and amputated tissue, in both nymphs and adults.

The concordant reduction of both ATAC-seq accessibility and H3K27ac and H3K4me1 histone marks indicates a massive loss of active regulatory elements. The fact that the same response occurs in regeneration-competent nymphs and regeneration-incompetent adults suggests that this genome-wide enhancer shutdown is associated with injury, regardless of whether tissues engage in regeneration. The molecular events that trigger regeneration should emerge within or after this injury-associated state.

### Comparative transcriptomics identify GbZelda as a candidate regulator of regeneration

To identify transcriptional responses associated with regenerative competence, we compared expression dynamics following leg amputation in regenerating stage 7 nymphs and non-regenerating adults. This comparison revealed a relatively small group of genes with distinct responses between the two developmental contexts (Fig. 3A,B; Supplementary Fig. 6). Several were noteworthy for their previously reported links to development or regeneration: Myoglanin, a recently characterized nymphal development factor in crickets, was upregulated (Kawamoto et al., 2025), whereas SmydA methyltransferase family members were downregulated, paralleling their response shortly after limb amputation in the Chinese mitten crab (Wang et al., 2022). Prominent changes also included upregulation of Ets21C, a transcription factor required to sustain a pro-regenerative transcriptional program during Drosophila imaginal disc regeneration (Worley et al., 2022). Among the most strongly induced genes was the *G. bimaculatus* ortholog of the *Drosophila* zinc-finger transcriptional activator Zelda, hereafter referred to as GbZelda (Fig. 3A).

**Figure 3.**
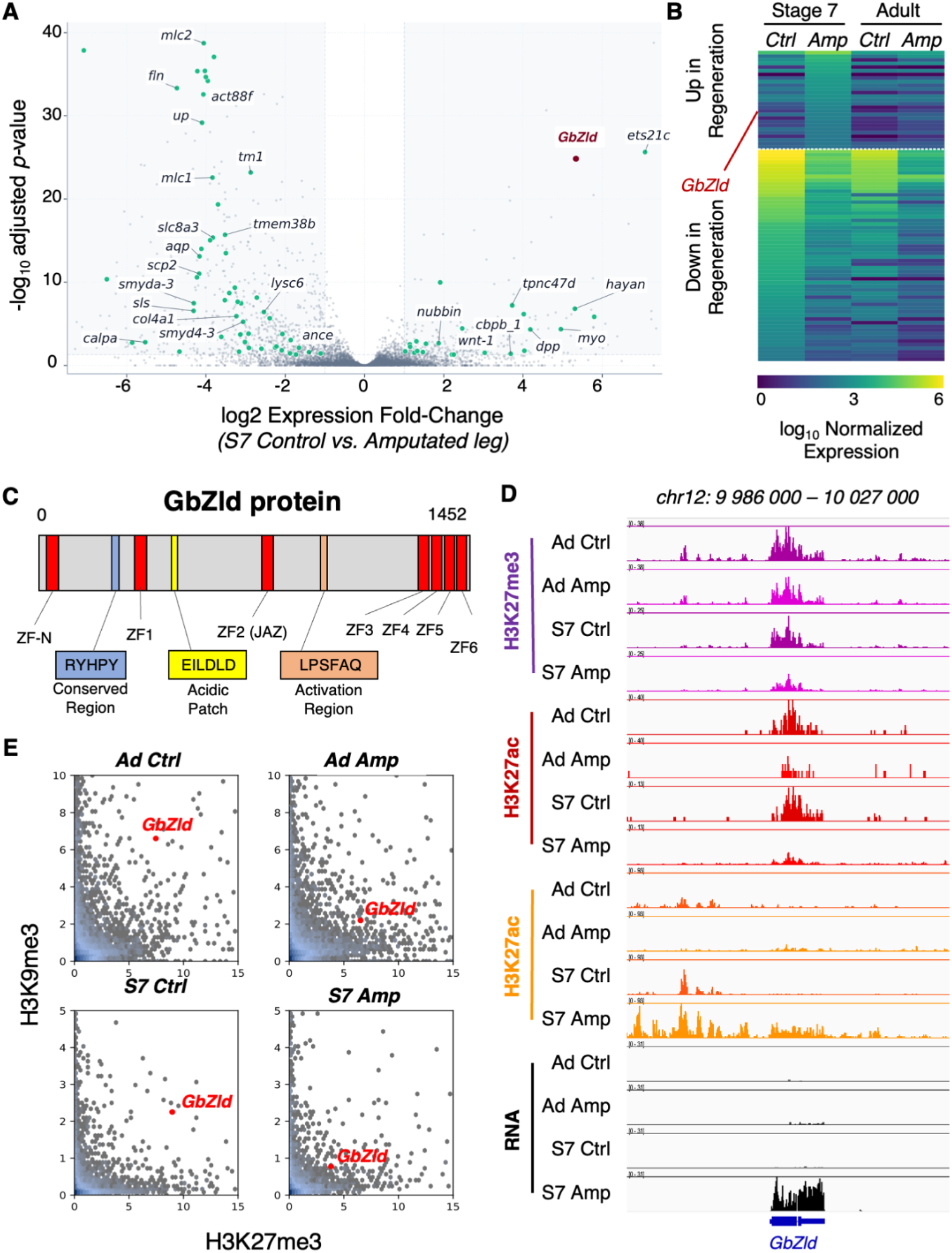
Regeneration-specific upregulation of GbZelda, associated with release of a double heterochromatin lock. (A) Differential expression analysis comparing stage 7 control and amputated legs, highlighting GbZelda among the genes that are upregulated during regeneration. (B) Expression profiles of genes induced or repressed specifically during regeneration. (C) Domain organization of the predicted GbZelda protein. (D) Chromatin profiles across the GbZelda locus. (E) Distribution of H3K27me3 and H3K9me3 histone modifications across gene bodies, highlighting GbZelda.

The predicted GbZelda protein contains the characteristic organization of pancrustacean Zelda proteins, including seven zinc-finger domains together with conserved regions associated with transcriptional activation (Fig. 3C)(Supplementary Figure 6). The six most C-terminal zinc fingers are strongly conserved among insect Zelda proteins, suggesting that the protein might recognize related sequences in different species. In *Drosophila*, Zelda acts as a pioneer transcription factor promoting enhancer activation during zygotic genome activation (Harrison et al., 2011; Zaret and Mango, 2016; Calderon et al., 2022; Brennan et al., 2023). More recently, Zelda has also been implicated in the response to genetically induced tissue damage in *Drosophila* imaginal discs (Harris et al., 2020; Bose et al., 2025). The selective induction of GbZelda in the context of a genome-wide enhancer shutdown suggests it could also play a role as a pioneer transcription factor during appendage regeneration in crickets.

### A double chromatin lock on the *GbZelda* locus is selectively released in regenerating tissue

GbZelda expression increases strongly in amputated stage 7 legs, whereas the gene remains largely silent in adult amputated tissue (Fig. 3D). This difference is particularly striking given the observation that both tissues display the same broad loss of enhancer-associated chromatin. We therefore asked how GbZelda expression was permitted in the amputated legs of stage 7 nymphs, but not in adults. In particular, we reasoned that the selective induction of GbZelda must occur despite the genome-wide enhancer shutdown observed in both tissues, and therefore might be associated with locus-specific chromatin profiles. To address this question, we examined the chromatin state of the GbZelda locus in greater detail (Fig. 3D). In intact leg tissue, the GbZelda locus displays an unusual combination of H3K27me3 and H3K9me3 (Fig 3D). Indeed, the coexistence of these two marks is noteworthy because H3K27me3 and H3K9me3 are generally associated with distinct repressive chromatin environments (Janssen and Lorincz, 2022). H3K27me3 is classically associated with Polycomb-mediated repression of developmental genes, whereas H3K9me3 is a hallmark of heterochromatin. The observation that both marks occupy GbZelda in intact leg tissue suggests that expression of this developmental regulator is subject to unusually stringent control. This combination of chromatin marks is also unusual in the cricket leg epigenome: GbZelda stood out among the small number of loci enriched for both repressive modifications (Fig 3E).

Next we compared these two marks at the GbZelda locus before and after leg amputation. H3K9me3 is strongly reduced after injury in both stage 7 nymphs and adult legs (Fig 3D, E). In contrast, H3K27me3 is substantially reduced at the GbZelda locus in amputated stage 7 legs, where GbZelda is expressed, but persists in adult amputated legs, where GbZelda remains silent. Moreover, H3K27ac, which is broadly depleted after amputation across the genome, shows a local gain upstream of GbZelda in stage 7 amputated legs. The locus therefore provides a particularly clear example of the difference between the global injury response and the selective chromatin changes associated with regeneration. Combinations of H3K27me3 and H3K9me3 have been described in other contexts, where they are required for repression of target genes (Torres-Campana et al., 2022; Akkouche et al., 2025). We therefore propose that the GbZelda locus is constrained by a dual repressive chromatin state in intact leg tissue, which becomes released specifically in the regenerating legs of nymphs.

### Recovery of enhancers enriched for Zelda binding motifs

To explore whether the initial genome-wide enhancer shutdown could be followed by a selective reopening of regulatory elements mediated by GbZelda, we extended the ATAC-seq profiling to eight days after amputation, when stage 8 regenerating legs have undergone significant growth (Fig 4A).

**Figure 4.**
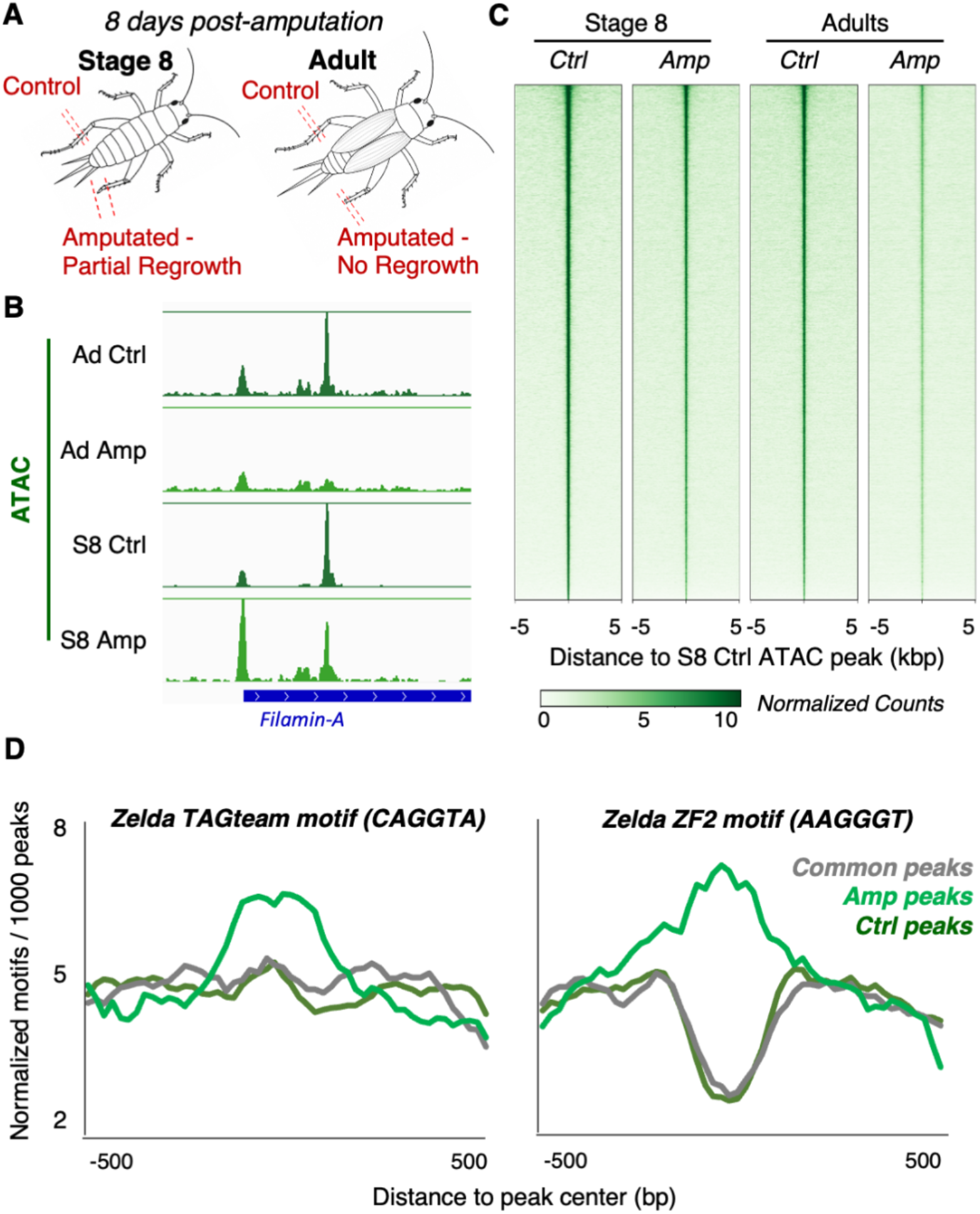
Partial chromatin reopening and Zelda motif enrichment associated with leg regeneration. (A) Experimental design for ATAC-seq at 8 days post-amputation. (B) Example ATAC-seq profiles in adult and stage 8 tissues. (C) Aggregate ATAC-seq profiles around stage 8 control peaks in control and amputated tissues. (D) Enrichment of Zelda binding motifs around common, control-enriched and amputation-enriched peaks.

At this later time point, the chromatin accessibility landscape still appears flatter and less structured in amputated legs compared with controls, but some peaks have started to re-emerge, most prominently in stage 8 regenerating legs (Fig 4B, C). Importantly, all the ATAC-seq peaks in these samples correspond to peaks that we also find in the non-amputated controls; we detect virtually no peaks that are specific to amputated/regenerating legs.

The extent of chromatin re-opening varies between peaks. To assess whether this might be associated with the action of GbZelda, we classified peaks into ‘High’, ‘Mid’ and ‘Low’ categories, depending on the extent of recovery of the ATAC-seq signal. In Drosophila, Zelda recognizes the TAGteam motif (CAGGTA) through its four C-terminal zinc fingers, ZnF3–6 (Hamm et al., 2015), and the recently identified AAGGGT motif through its second zinc finger, ZnF2 (Torres-Zelada et al., 2025). We find that the Zelda binding motifs CAGGTA and AAGGGT are enriched specifically in the ‘High’ recovery peaks. The enrichment of the latter motif is particularly striking, as the motif appears to be specifically depleted at the ‘Mid’ and ‘Low’ recovery peaks.

### Genome-wide chromatin remodelling is involved in leg regeneration in a distantly related arthropod species

To test whether the chromatin dynamics that we identified in the cricket are shared with regenerating legs in other arthropod species, we extended this analysis to the crustacean *Parhyale hawaiensis*, an experimental system that has already been used to study leg regeneration (Konstantinides and Averof 2014, Alwes et al 2016, Sinigaglia et al. 2022). Unlike crickets, *Parhyale* are able to regenerate their legs throughout their lifetime, from juvenile to adult stages. To assess changes in chromatin accessibility that occur during the early stages of leg regeneration, we performed ATAC-seq on uninjured *Parhyale* legs and on amputated leg stumps collected 1 day post amputation (dpa). We chose 1 day post amputation because the pace of leg regeneration in *Parhyale* is more rapid than in crickets (nearly complete regeneration within 10 days post injury; Cevrim et al. 2025).

ATAC-seq in uninjured *Parhyale* legs reveals a complex landscape of chromatin accessibility, in which a large proportion of reads fall in a small number of discrete peaks across the genome (average genome-wide FRiP scores 0.2 to 0.3); these peaks include the promoter regions of most genes and putative enhancers (Forbes et al. 2026). In contrast, the ATAC-seq profiles of leg stumps at 1 day post amputation show a much flatter landscape, with much fewer and shorter peaks (Fig 5B,C), which remind us of the profiles observed 3 days post amputation in crickets. These flat profiles are evident across the entire genome; less than 1% of genomic regions have a FRiP score greater than 0.2 (Fig 5D).

**Figure 5.**
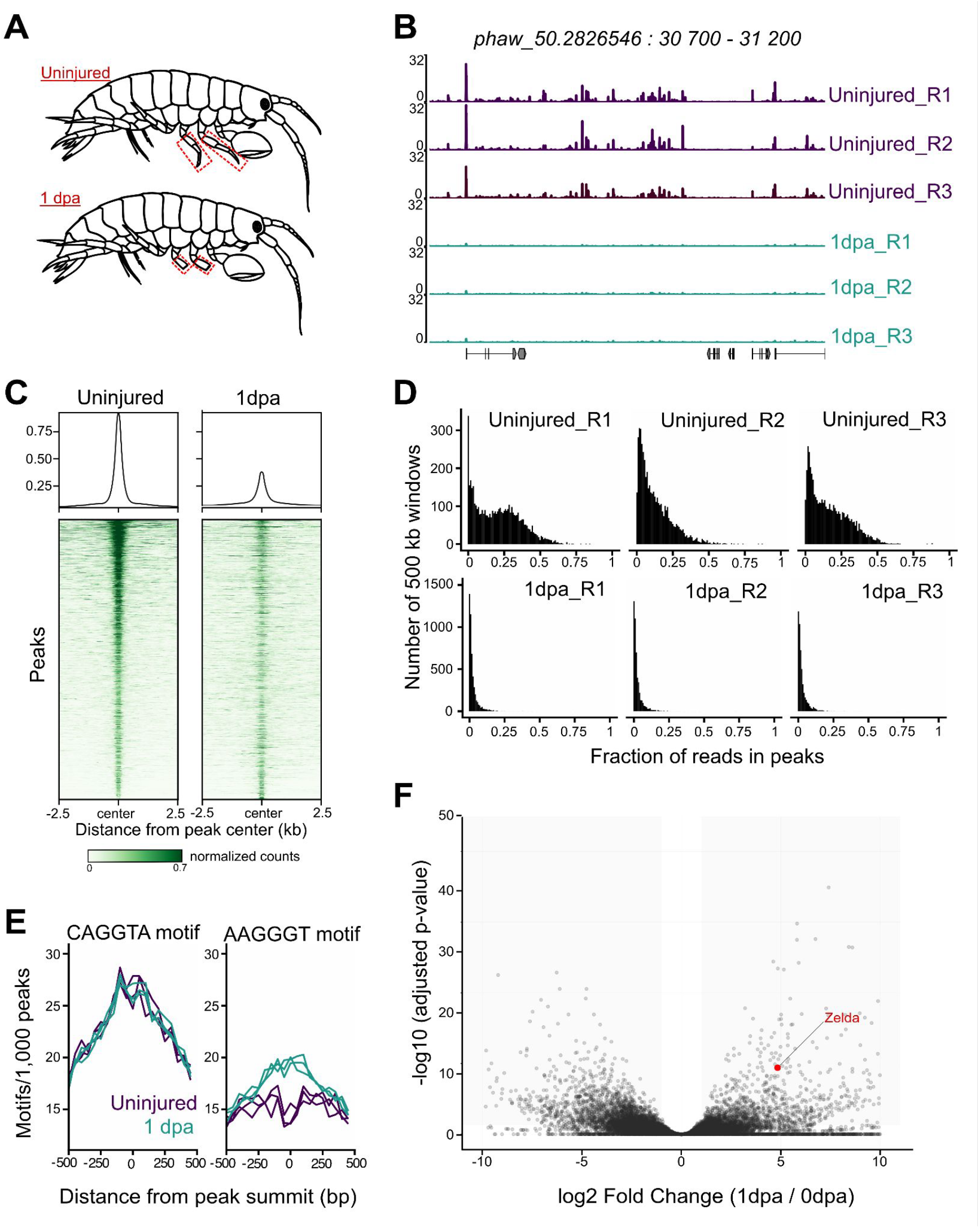
Chromatin accessibility profiles and Zelda enrichment in regenerating *Parhyale* legs. (A) Illustration of leg samples collected from uninjured and amputated (1 day post injury, 1 dpa) *Parhyale* legs. (B) Genome browser profiles across a 500 kb region, illustrating the loss of discrete ATAC-seq peaks 1 day post amputation (3 replicates). (C) Enrichment of ATAC-seq signals around ATACseq peaks identified in uninjured and 1 dpa legs. (D) Distribution of FRiP scores across the genome (measured on 500 kb genomic fragments) in uninjured and 1 dpa legs. In uninjured legs, 20 to 50% of genomic fragments have FRiP >0.2; in contrast, at 1 dpa, fewer than 1% of fragments have FRiP >0.2. (E) Enrichment of the Zelda binding motifs CAGGTA and AAGGGT in the vicinity of open chromatin peaks (+/-500 bp from the peak summit) in uninjured and 1 dpa legs (3 replicates). (F) Differential expression analysis comparing uninjured (0dpa) and amputated legs (1dpa), highlighting Zelda among the genes that are upregulated at 1 dpa (the plot has been cropped, for the entire plot see Suppl. figure 7).

We find that the two Zelda-binding motifs are differentially enriched in the ATACseq peaks in uninjured and newly amputated legs: the CAGGTA motif is enriched in peaks both before and after amputation, whereas the AAGGGT motif is specifically enriched in the peaks that are detected after amputation (Fig 5E). As in crickets, we find that the orthologue of Zelda (MSTRG.29072) is upregulated during the early phases of leg regeneration in *Parhyale*, starting from 12 hours post amputation (Fig 5F and Supplementary Fig. 7).

These results suggest that within a day from leg amputation, cells in the injured leg stump drastically re-model their chromatin. The profiles of chromatin accessibility become flatter on a genome-wide scale, with most peaks of open chromatin closing down. Within this flattened landscape, a subset of promoters, which are enriched in Zelda binding motifs, remain accessible, coincident with the upregulation of Zelda expression.

## DISCUSSION

The central finding of our study is that appendage regeneration in arthropods involves a reproducible sequence of chromatin remodelling events. In *G. bimaculatus*, limb amputation is followed first by a striking genome-wide shutdown of enhancer-associated chromatin, with loss of discrete accessible regions and depletion of H3K27ac and H3K4me1 histone marks. In regeneration-competent nymph legs, this broad response is followed by the selective de-repression of GbZelda expression and a re-opening of enhancers that are enriched in Zelda binding motifs. In *P. hawaiensis* we observe a similar succession of enhancer shutdown and Zelda upregulation during early stages of regeneration. These results suggest that regeneration proceeds through remodelling of chromatin on a genome-wide scale, rather than by regulating a limited set of regeneration-specific enhancers. We propose that these genome-wide changes serve to erase some of the epigenetic memory that lock cells into terminally differentiated cell states, allowing the plasticity needed to engage in re-building the leg.

The developmental decline of regenerative abilities in crickets, from early nymphal stages to adults, provide a useful framework for distinguishing changes that follow leg amputation in all circumstances, from ones associated more specifically with tissues that proceed toward regeneration. The genome-wide enhancer shutdown, which occurs in both stage 7 nymphs and adults, cannot itself explain the different regenerative abilities of nymphs and adults. Instead, the subsequent events, involving the selective re-opening of chromatin mediated by Zelda, are likely to contribute to the different regenerative outcomes at these stages. The similarities we observe in crickets and crustaceans suggest these events are ancient and widely conserved aspects of the regenerative process in pancrustaceans, and possibly more widely.

The most unexpected feature of this response is the extent of the early enhancer shutdown observed in crickets and crustaceans. In many regenerative systems, injury is associated with the activation of developmental or regeneration-responsive enhancers (Kang et al., 2016; Wang et al., 2020; Harris, 2022; Kawaguchi et al., 2024). Here, by contrast, the earliest regulatory events that we observe after amputation are characterized by a widespread loss of chromatin accessibility. The observation that these changes are accompanied by depletion of H3K27ac and H3K4me1 histone marks in crickets, suggests there is a coordinated mechanism of enhancer shutdown. This state of genome-wide enhancer shutdown appears to be the starting point for a selective genome re-activation, mediated by Zelda, in the regenerating legs of crickets and crustaceans.

Zelda is a well-studied pioneer transcription factor in *Drosophila*, where it plays a key role in the transcriptional activation of the zygotic genome (Harrison et al., 2011; Zaret and Mango, 2016; Calderon et al., 2022; Brennan et al., 2023) and in exit from regeneration following damage-induced tissue regrowth (Bose et al., 2025). Our results suggest a similar role in selective genome activation during leg regeneration in crickets and crustaceans. Although we have not directly tested the role of Zelda through functional experiments, the discovery that Zelba binding motifs are enriched in similar patterns in arthropods separated by more than 500 million years of divergent evolution suggests that Zelda plays an important and deeply conserved role during leg regeneration in arthropods.

The chromatin state of the GbZelda locus in crickets provides a further clue to its role in regeneration. In intact cricket legs, GbZelda is unusually marked by both H3K27me3 and H3K9me3, a combination of histone marks that is rare among genes in our dataset. Following amputation, H3K9me3 is reduced in both nymphal and adult tissues, and H3K27me3 is selectively reduced in regenerating nymphs, coincident with GbZelda activation. These results suggest that expressing Zelda requires lifting two layers of repression, one which is lifted in all amputated legs and a second that is relieved only in regeneration-competent nymphal legs. This dual repression mechanism likely provides a tight checkpoint during the early stages of leg regeneration.

Taken together, our findings support a model in which appendage regeneration is associated with a two-phase chromatin remodelling that is conserved across pancrustaceans. Tissue loss first triggers a genome-wide shutdown of enhancer-associated chromatin. In regeneration-competent tissues this shutdown is accompanied by a selective reopening of specific *cis-*regulatory elements, mediated by the pioneer transcription factor Zelda. This mechanism of regulated genome shutdown and re-opening, which has been conserved over hundreds of millions of years of evolution in pancrustaceans, raises interesting questions about whether and how such re-programming occurs in other regenerative organisms, across the animal kingdom.

## MATERIALS & METHODS

### Animal husbandry and cricket physiology assessment

All animal manipulations were performed under CO_2_ anesthesia. Gryllus bimaculatus crickets were maintained under controlled laboratory conditions. Females were individually placed in egg-laying containers for 6–10 h, and hatching dates were recorded to establish synchronized developmental cohorts. Molting events were monitored throughout development.

### Leg regeneration quantification assays

Regeneration experiments were performed by amputating the tibia at the midpoint of the third thoracic leg under CO_2_ anesthesia. Amputated individuals were returned to their rearing containers and monitored after subsequent molts. Regeneration was assessed by measuring regenerated tibia length and comparing regenerated legs with the contralateral intact legs.

To account for individual growth differences, regeneration efficiency was normalized to the growth of the non-amputated leg:

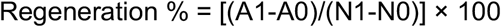

where N_1_ and N_0_ correspond to measurements of non-amputated legs after regeneration and at day 0, respectively, and A_1_ and A_0_ correspond to regenerated and initial amputated leg measurements. Measurements were performed 8 and 15 days post-amputation, following a molt to ensure comparable developmental stages.

### Assembly and scaffolding of the *Gryllus* genome

Approximately 10^6^ cells in 1 mL 1X PBS were fixed with 1% formaldehyde (Sigma-Aldrich, cat. F8775) for 10 minutes at RT with orbital agitation at 120 rpm. Formaldehyde was quenched with 200 mM glycine for 5 minutes at RT at 120 rpm. Cells were washed with cold 1X PBS at 300 g for 5 minutes. The pellet was flash frozen in liquid nitrogen and stored at -80°C. The pellet was thawed on ice, resuspended in cold H_2_O supplemented with an anti-protease mix (Roche, cat. 11836170001) and transferred to a Precellys VK05 tube, and lysed for 3 × 30 s at 6,800 rpm. Approximately 10^6^ cells were processed for Hi-C using the Arima Hi-C+ kit (Arima Genomics, cat. A410079) following the manufacturer’s instructions.

De novo genome assembly was generated using Oxford Nanopore Technology long reads with Flye (v2.9.3-b1797). The assembly was polished using publicly available Illumina short reads from SRA (DRX261892, DRX261893, DRX261894, DRX261895, DRX261896, and DRX261897) with HyPo (v1.0.3). Genome scaffolding was performed using InstaGRAAL (v0.1.6), followed by additional polishing with HyPo. The scaffolded genome was manually curated to properly separate chromosomes. Approximately 300 kb of sequence remained unscaffolded and was retained as “unscaffolded_contig” in the final reference genome.

### RNA sequencing and differential expression analysis

For RNA-seq experiments, the third thoracic leg (T3) was amputated at the midpoint of the tibia immediately after molting to induce regeneration. Regenerating stumps and corresponding contralateral intact legs were collected three days post-amputation. Each biological replicate consisted of pooled tissues from four individuals (two males and two females), with three biological replicates per condition.

Four sample types were analyzed: adult amputated legs, adult contralateral control legs, stage 7 amputated legs, and stage 7 contralateral control legs.

Total RNA was extracted from collected tissues using the RNeasy Mini Kit (Qiagen, #74104) according to the manufacturer’s instructions. Poly(A) mRNA libraries were prepared from total RNA using the Illumina Stranded mRNA Prep Ligation kit according to the manufacturer’s instructions. Sequencing was performed on an Illumina NextSeq 2000 platform using a 100-cycle kit to generate 50 bp paired-end reads (PE50). Sequencing yielded an average of 30 million reads per sample across the 12 multiplexed samples. The raw sequencing reads were pre-trimmed of adapters by the sequencing facility prior to delivery. Raw sequencing data were processed using the Gryllus bimaculatus reference genome described above.

Raw sequencing reads were aligned to the Gryllus bimaculatus reference genome using HISAT2 (v2.2.2) with customized splice alignment parameters (canonical splice penalty set to 0, non-canonical splice penalty to 3, minimum intron length of 20 bp, and maximum intron length of 500,000 bp, with transcriptome assembly reporting enabled via --dta). Alignments were filtered to retain reads mapped in a proper pair with MAPQ ≥ 30 using Samtools (v1.8). RNA-seq downstream data analysis was performed in R (v4.3) using Bioconductor packages. Differential expression analysis was performed using DESeq2 with independent hypothesis weighting (IHW) for multiple testing correction.

To isolate the Stage 7 nymph-specific regeneration program, we implemented a two-step pipeline combining relative differential expression with absolute transcription volume filters:

1. Relative Interaction Modeling: We fitted a generalized linear model (GLM) with a 2×2 factorial interaction design (∼ Stage + Treatment + Stage:Treatment). A Likelihood Ratio Test (LRT) comparing the full model against a reduced model (∼ Stage + Treatment) identified genes whose relative transcriptional response to amputation significantly diverged between developmental stages (padj < 0.15 and absolute |log_2_FoldChange| ≥ 1.5, yielding 82 relative interaction genes).
2. Absolute Transcription Volume Rescue: Because log-ratio models can overlook high-abundance regulators with massive absolute pool size drops but similar relative ratios in both stages, we applied a three-fold biological volume rescue filter. Genes were rescued if they exhibited: (a) baseline significance in nymphs (padj < 0.05 and |log_2_FoldChange| ≥ 1.5), (b) an absolute change in normalized counts of ≥ 500 reads upon amputation in nymphs, and (c) a nymphal absolute change at least 3-fold larger than the adult absolute change (|ΔReads_Nymph| ≥ 3 × |ΔReads_Adult|, recovering 62 high-abundance regulators).

Combining the 82 base interaction genes and 62 rescued genes created a final 144-gene Hybrid Rescued Core, which was utilized for downstream functional enrichment and visualization. Differential expression results were visualized using ggplot2 (v3.4.0), ggrepel (v0.9.1), and dplyr (v1.0.10).

### Epigenomic profiling

Chromatin accessibility and histone modification profiles were assessed using CUT&RUN (in ovarian tissues), CUT&Tag (in regenerating and control leg tissues), and ATAC-seq (in both ovarian and regenerating leg tissues).

For regeneration experiments, the third thoracic leg (T3) was amputated at the midpoint of the tibia immediately after molting. Regenerating stumps and matched contralateral intact legs were collected three days post-amputation. Biological replicates consisted of pooled tissues from mixed-sex individuals. For ovarian samples, three females were collected per biological replicate, and experiments were performed in duplicate.

### CUT&RUN chromatin profiling

CUT&RUN experiments were performed on ovarian tissues. Tissues were isolated and immobilized on Concanavalin A-coated magnetic beads before antibody incubation. Chromatin profiling was performed using antibodies against H3K27me3 (Cell Signaling Technology, #9733), H3K9me3 (Abcam, ab8898), and H3K4me3 (Active Motif, #39159). IgG controls were included.

Following antibody binding, chromatin cleavage was performed using protein A–MNase (pA-MNase). Released DNA fragments were purified using AMPure XP beads (Beckman Coulter, A63881) and quantified using a Qubit dsDNA High Sensitivity assay (Thermo Fisher Scientific). Cleaved chromatin libraries were prepared using the Diagenode CUT&RUN library preparation kit according to the manufacturer’s protocol. Sequencing was performed on an Illumina NextSeq 2000 platform to generate 50 bp paired-end reads, yielding approximately 50 million reads per sample across 8 multiplexed histone mark samples (four marks in duplicate) and IgG controls.

### CUT&Tag chromatin profiling during regeneration

CUT&Tag was performed on regenerating and control leg tissues. Nuclei were isolated from pooled tissues and incubated with antibodies targeting histone modifications associated with repressive chromatin states: H3K9me3 (Abcam, ab8898) and H3K27me3 (Cell Signaling Technology, #9733). IgG controls (Merck, NI01) were included.

Chromatin tagmentation was performed using pA-Tn5 transposase (Diagenode, C01070001). DNA fragments were purified using the MinElute PCR Purification Kit (Qiagen, #28204). Libraries were amplified using NEBNext HiFi 2× PCR Master Mix (New England Biolabs, M0541L) and purified using AMPure XP beads (Beckman Coulter, A63881). Library quality was assessed using an Agilent Bioanalyzer 2100 High Sensitivity DNA assay, and sequencing was performed on an Illumina NextSeq2000 platform (paired-end 50 bp reads) at the Institut Curie Next Generation Sequencing Platform.

### ATAC-seq chromatin accessibility profiling

ATAC-seq was performed on ovarian and regenerating leg tissues. Nuclei were isolated and processed using the Active Motif ATAC-seq Kit (#53157) according to the manufacturer’s instructions with minor modifications. For regeneration experiments, amputated stumps and matched contralateral control legs were compared as “regeneration” and “non-regenerative” conditions, respectively. ATAC-seq experiments in regenerating leg tissue were performed at 3 days post-amputation and were additionally performed at 8 days post-amputation to assess the later chromatin response. Sequencing was performed on an Illumina NextSeq 2000 platform to generate 50 bp paired-end reads, yielding approximately 30 million reads per sample across 5 processed samples.

### Epigenomic data processing and peak calling

Raw sequencing files were quality-checked using FastQC (v0.74), and no downstream adapter trimming was performed since adapters were pre-trimmed by the sequencing facility prior to delivery. Paired-end reads were aligned to the Gryllus bimaculatus reference genome using Bowtie2 (v2.5.5) for ATAC-seq, CUT&Tag, and CUT&RUN, with parameters --very-sensitive -X 2000 --no-mixed --no-discordant --dovetail (for ATAC-seq) or --very-sensitive -I 10 -X 700 --dovetail --no-mixed --no-discordant (for CUT&RUN and CUT&Tag). For ATAC-seq, PCR duplicates were marked using Picard MarkDuplicates (v3.1.1.0) and filtered using deepTools alignmentSieve (v3.5.4) in paired-end mode with MAPQ ≥ 20, --ignoreDuplicates, and exclusion of bitwise flag 1024. For CUT&RUN and CUT&Tag, PCR duplicates were marked using Picard MarkDuplicates (v3.1.1.0) and physically removed, followed by filtration using Samtools (v1.22) for properly paired reads (SAM flag 2) with MAPQ ≥ 30 (for CUT&RUN and CUT&Tag H3K27me3) or MAPQ ≥ 1 (for CUT&Tag H3K9me3). For CUT&RUN and CUT&Tag, fragments ≤ 700 bp were sieved using deepTools alignmentSieve (v3.5.4). For ATAC-seq, alignmentSieve was used to shift fragment ends as appropriate.

Open chromatin regions (ATAC-seq) and histone modification enrichment sites (CUT&RUN and CUT&Tag) were identified using peak-calling software. MACS2 (v2.2.9.1) was used for narrow peaks (H3K4me3, H3K27ac) and open chromatin regions with a strict q-value threshold of < 0.05 (effective genome size: 1,660,000,000 bp). Broad marks (H3K27me3, H3K9me3) for both CUT&RUN and CUT&Tag were evaluated using SEACR (v1.3) run in stringent mode. Genomic tracks (BigWig/BedGraph format) were generated using deepTools bamCoverage (v3.5.4) normalized using 1× sequencing depth (RPGC, effective genome size: 1,660,000,000 bp) with exclusion of the unscaffolded_DNA region for normalization. For CUT&Tag and ATAC-seq tracks, reads were centered. For ATAC-seq, tracks were normalized using Counts Per Million (CPM). Differential enrichment analysis across conditions was computed using DESeq2 (v1.40.2 / v2.11.40.8). RNA-seq, ATAC-seq and chromatin mark enrichment data were visualized as heatmaps, in which each cell represents an enrichment result, each line represents a promoter or ATAC peaks, and color indicates the value according to the accompanying color scale. This format allows patterns across categories and conditions to be compared visually. To compute chromatin enrichment profiles over gene bodies, deepTools computeMatrix (v3.5.4) was executed in scale-regions mode which was used later for scatter plot generation, and the overall heatmaps were visualized using plotHeatmap (v3.5.4).

### ATAC-seq profiling and analyses in *Parhyale* legs

*Parhyale hawaiensis* of the inbred line Chicago-F (Kao et al. 2016) were cultured in artificial sea water using standard methods (Browne et al. 2005). ATAC-seq was performed on uninjured and one day post amputation T4 and T5 *Parhyale* legs. Uninjured samples were previously published (Forbes et al. 2026). For regenerating samples, animals were anaesthetised in 0.02% clove oil in artificial seawater and legs were amputated at the distal end of the merus 2-5 days post molting. Regenerating tissue was collected by cutting the proximal end of the merus. Nuclei were isolated on a cold block by opening the cuticle and removing the tissue using a surgical knife (Fine Science Tools #10316-14). Cuticle and tissue were incubated separately in 125 µl of a lysis buffer (10 mM Tris-HCl pH 7.4, 10 mM NaCl, 3mM MgCl2, 1% BSA, 0.1% Igepal CA-630, 0.1% Tween-20, 0.01% digitonin) and passed 25 times through a P200 micropipette tip. Supernatant from the cuticle portion was combined with the lysed tissue portion to remove contamination of large cuticle pieces. An equal volume of wash buffer (10 mM Tris-HCl pH 7.4; 10 mM NaCl; 3mM MgCl2; 1% BSA; 0.1% Tween-20) was then added and the lysate was passed through a 10 µm pluriStrainer cell strainer (pluriSelect #43-10010) to remove any remaining cuticle. 8 to 17 thousand nuclei were recovered per sample. Tagmentation was performed in 50 µl tagmentation mix, containing tagmentation buffer (Diagenode #C01019043) and 2.5 µl Tn5 loaded transposase (Diagenode #C01070012) at 37°C for 30 minutes, shaking at 300 RPM. Paired-end 76 bp sequencing was firstly performed on Illumina NextSeq 500 sequencer followed by deeper sequencing on Illumina NovaSeq. NovaSeq reads were trimmed from 150 bp to 76 bp and Nextera adapters removed using cutadapt (version 4.4) with a minimum overlap of 5 and quality trimming of 20. Adapter sequences were removed from NextSeq reads by the sequencing facility. Mapping and peak calling were performed as previously described (Forbes et al. 2026).

Reads were mapped to the genome and mitochondrial, duplicate and low quality mapping reads were removed as described in Forbes et al. 2026. Alignments were shifted using alignmentSieve ATACshift from deepTools (v 4; Ramírez et al., 2016). Peaks were then called using MACS2 (v 2.2.7.1; Zhang et al., 2008) in paired-end mode, with an effective genome size of 2.8 Gb and q-value threshold of < 0.05. Genome browser plots were generated by pyGenomeTracks (v 3.9, Lopez-Delisle et al., 2021) using the MACS2 bedGraph files, normalised by fragments per million.

The fraction of reads in peaks (FRiP) was calculated as the number of reads overlapping with the MACS2-defined peaks divided by the total number of reads. To determine whether differences in FRiP could be attributed to sequencing depth, all samples were downsampled to 10 million reads (5 million fragments), after which peaks were re-called using the same MACS2 parameters (Supplementary Table 1).

To visualise the difference in ATAC-seq signal across all peaks (Fig. 5C), MACS2 bedgraph files from the downsampled datasets were merged across replicates using bedtools unionbedg (v 2.31.1; Quinlan and Hall, 2010) and normalised to fragments per million. The normalised bedGraph files were converted to bigWig format by bedGraphToBigWig (v 2.10; Kent et al., 2010). A signal matrix was then generated using deepTools computeMatrix, using a merged set of all called peaks as the reference regions. The reference-point was set at centre, with a bin size of 10 and and a flanking region of +/-2.5 Kb. Heatmaps were generated using deepTools plotHeatmap, with the peaks ranked by their mean in the uninjured.

To assess the distribution of FRiP scores across the genome (Fig. 5D), the genome scaffolds were split into 500 kb windows, and windows less than 500 kb were removed from the analysis. Using the downsampled datasets, paired-end fragments were counted within each window using featureCounts (Rsubread version 2.24.0) (Liao et al.,2014). Regional FRiP was calculated as the number of fragments overlapping the previously defined MACS2 called peak within each window, divided by the total number of fragments in that window. Only windows with at least 50 fragments were plotted.

Enrichment of the two Zelda binding motifs (CAGGTA and AAGGGT, Fig. 5E) was assessed across MACS2 defined peaks from each sample. Peaks overlapping with naked DNA peaks (called with the same MACS2 parameters as for the uninjured and 1 dpa samples) were excluded prior to the analysis to minimise bias of the sequence preference of the Tn5 transposase. The naked DNA dataset was previously published in Forbes et al. 2026. A region +/-500 bp of the peak summit was searched for presence of motifs and a bin size of 50 bp was used.

Gene expression analysis analysis of *Parhyale* legs (Fig. 5F) was done using the published regeneration bulk RNA-seq dataset (Sinigaglia et al. 2022). We used the 11 single-leg samples collected at 0 and 1 dpa (GEO accession numbers GSM5885168, GSM5885169, GSM5885170, GSM5885171, GSM5885172, GSM5885178, GSM5885179, GSM5885180, GSM5885181, GSM5885182 and GSM5885183); in most cases pairs of samples originated from the same individual. We used DESeq2 v1.50.2 (Love et al., 2014) on raw counts for identifying genes differentially between 0 and 1 dpa. We noticed a very strong correlation in samples coming from the same individual. To avoid individual effects, we pooled the samples coming from the same individuals using the DESeq2 function collapseReplicates. We also performed this analysis without pooling, with very similar results.

## ACKNOWLEDGEMENTS

We thank Dr. Mélody Nicolau, Lisa Germain and Dr. Sophie Barral for help with cricket husbandry. We thank Dr. Benjamin Loppin, Dr. R. Dubruille, Dr. Antoine Molaro and Dr. Guillaume Martinez for feedback throughout the project, as well as the entire Epigenetics and Chromatin team at IAB for helpful discussions. The research on crickets was supported by grants from Agence Nationale de la Recherche (ANR JCJC CRACK; ANR PRC Shade-of-Pal; ANR PRC SMMILE). ML was supported by a 4th year fellowship from Fondation pour la Recherche Médicale. The research on Parhyale was supported by a grant from the Fondation pour la Recherche Médicale (grant EQU202303016278). We thank the sequencing platforms GenomEast and ICGex, supported by Agence Nationale de la Recherche.

## AUTHOR CONTRIBUTIONS

ML conducted most of the cricket experiments. GF generated the ATAC-seq data from Parhyale. GF and MP analysed the Parhyale data. MMA and FC analysed the cricket epigenomics and transcriptomic data. EM generated CUT&Tag data. AA performed cricket regeneration tests. FG and RK assembled the cricket genome. AD and AP generated cricket Hi-C data. BP, CS and MAH generated cricket Nanopore data. MA and MP conceived and supervised the Parrhyale project. GO conceived and supervised the cricket project. ML, GF, MA, MP and GO wrote the manuscript.

