## Supplemental Figures for "Genome-wide chromatin reset at the onset of leg regeneration in pancrustaceans"

### Supplementary Figure 1

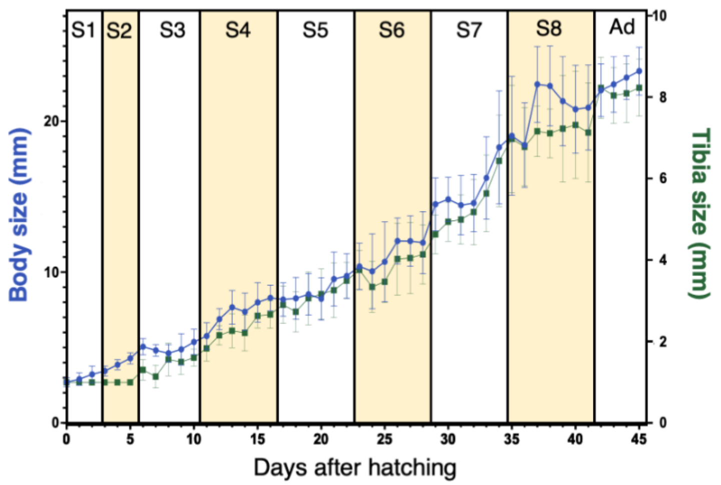


**Normal growth of whole *G.bimaculatus* individuals and tibias across nymphal development**

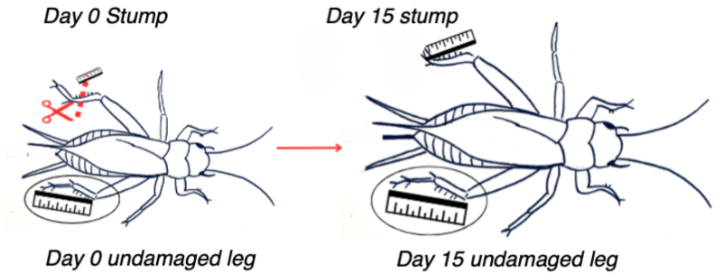


**Schematic representation of regeneration quantification experiment**

#### Supplementary Figure 2

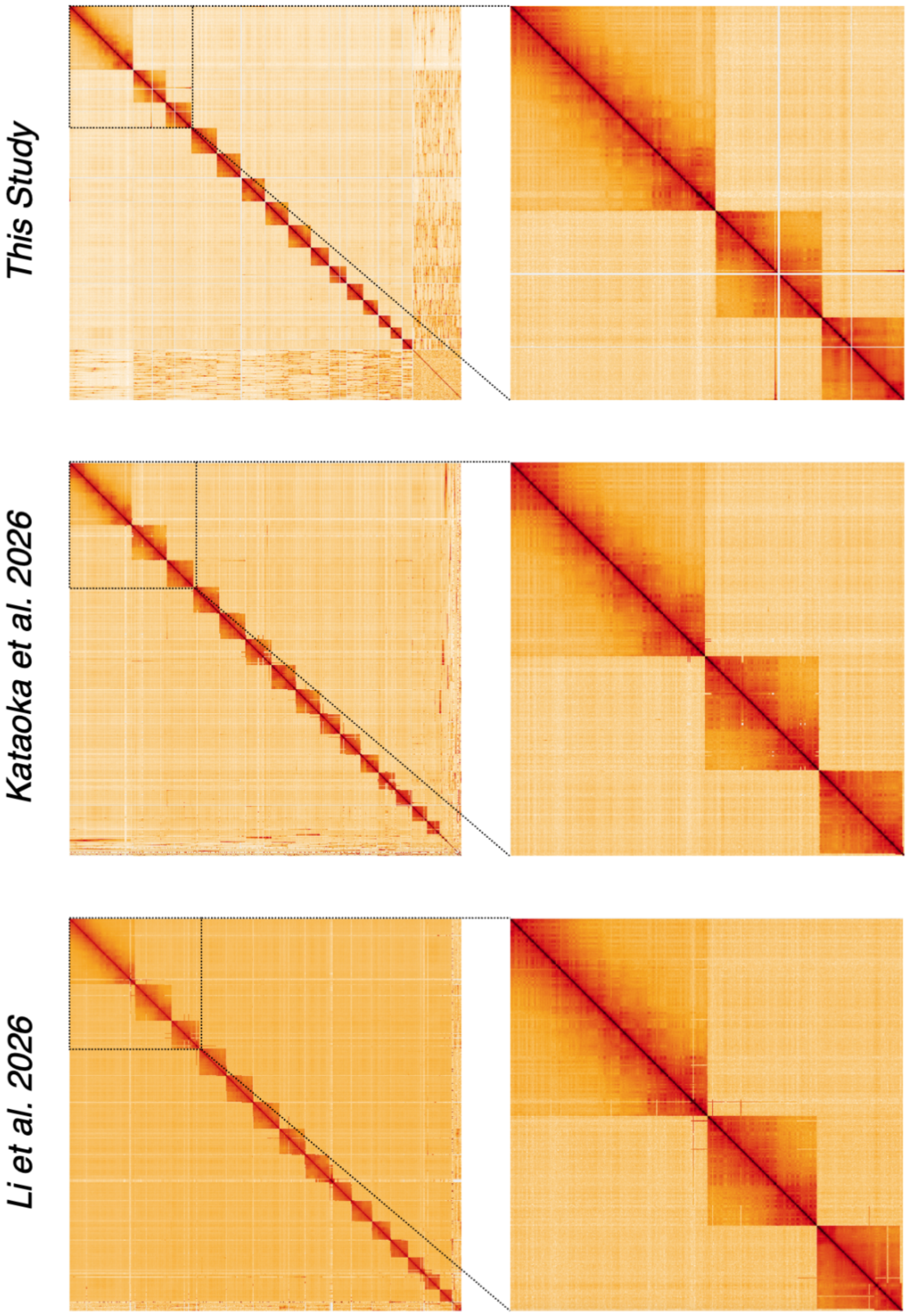


**Hi-C data mapping to 3 versions of the *G.bimaculatus* genome.**

### Supplementary Figure 3

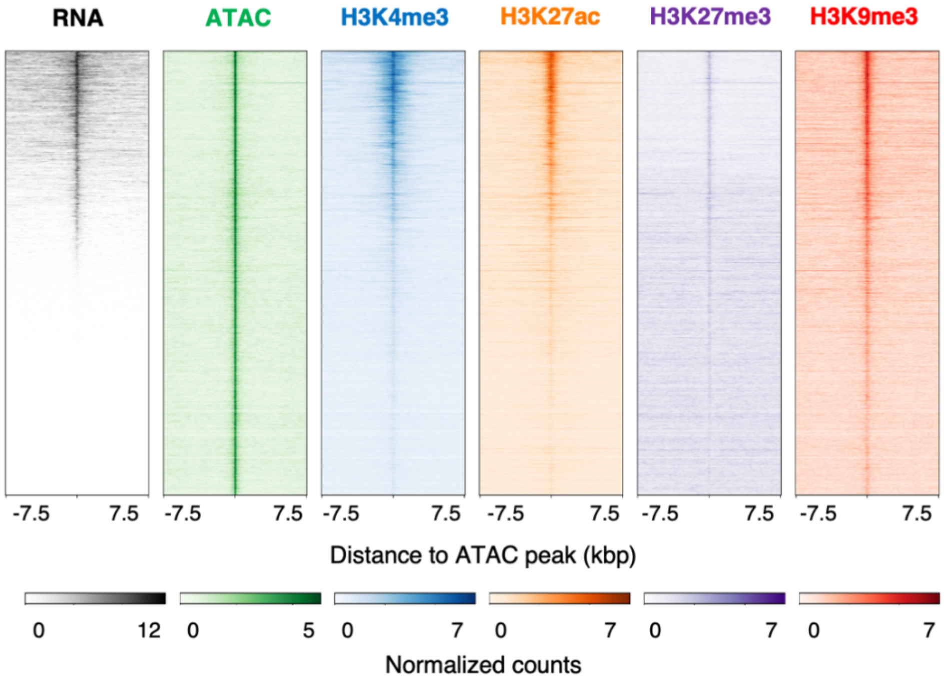


**RNA, ATAC signal and Histone modification distribution around Ovary ATAC-seq peaks.**

#### Supplementary Figure 4

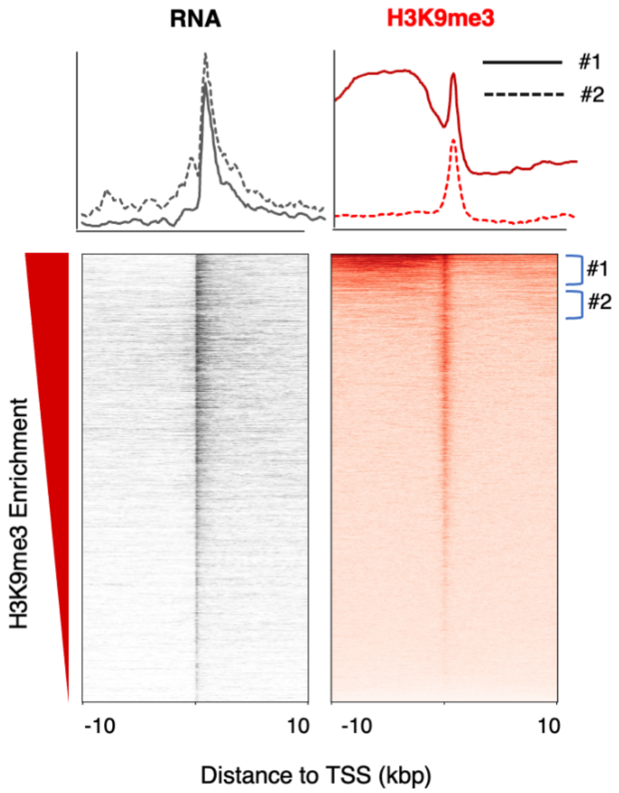


**H3K9me3 is enriched upstream of highly expressed genes where it limits transcription.**

### Supplementary Figure 5

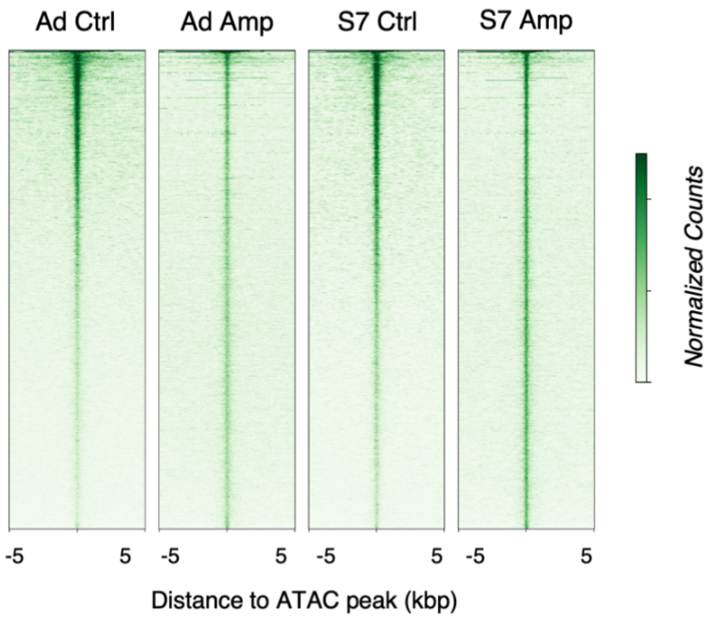


**ATAC signal at Regenerating Stage 7 ATAC peaks.**

### Supplementary Figure 6

```
                 ---------ZF3----------      ----------ZF4----------      ---------ZF5----------          ---------ZF6-----
Gryllus       HKCLDCDKVFNKACYLTQHNKSFHAGEKPFKCKRCGKRFPAEPLYAEHCSKH-AGDKPHKCSLCPKQFNHKTDLRRHI-CLHTGDKPFACGVCGKGFIRKDHMLKHCETHRKK
Drosophila    IKCLECDKEFTKNCYLTQHNKSFHSGEYPFRCQKCGKRFQSEDVYTTHLGRHRTQDKPHKCELCPKQFHHKTDLRRHVEAIHTGLKQHMCDICEKGFCRKDHLRKHLETHNRP
Blattella     IICIECDKVFNKACYLTQHNKSFHAGHKPYKCDRCGKRFTTECMYTEHLGKH-AGEKPHKCDACPKQFNHKTDLRRHM-CLHTGEKPFTCDICKKGFIRKDHMLKHCETHRRK
Zootermopsis  HKCVDCDKVFNKACYLTQHNKSFHSGHKPYKCDRCGKRFMIEMTYEQHLAKH-AGEKPYKCDVCPKQFNHKTDLRRHM-CLHTGMKPFQCNICSKGFIRKDHMLKHCETHRRK
Tribolium     YKCIDCDKTFNKACYLTQHNKTFHCGDKPFKCTRCGKRFSNEETHEEHASKH-AGDKPHKCDRCPKQFNHKTDLRRHM-CLHTGQKPFVCDTCGKGFIRKDHMLKHCETHFRK
Agrilus       HKCFDCDKMFNRACYLTQHNKTFHSGDKPFKCTRCGKRFSCEKQYQNHSTKH-AGDKPYKCDLCPKQFNHKTDLRRHM-CLHTGQKPFACDTCGKGFIRKDHMLKHCDTHSRR
Danaus        HKCFECDKLFNKVCYLTQHNKTFHSGAKPFKCDRCGKRFSDGVSYEGHYLKH-ADNKPFKCNECPKSFNHKTDLRRHM-CLHSGCKPFACDHCGKGFIRKDHMVKHFDTHIKK
Apis          HKCIECDKVFNKACYLTQHNKSFHSGDKPFKCNQCGKRFPLEYLHAEHLQKH-AGDKPYKCEICPKQFNHKTDLRRHL-CLHTGEKPYACDNCGKGFIRKDHMMKHLETHKKK
Nasonia       HKCIECDKVFNKACYLTQHNKSFHSGDKPFKCNMCGKRFPHEYLHAEHLQKH-AGDKPYKCEICPKQFNHKTDLRRHM-CLHTGEKPYACDKCGKGFIRKDHMMKHLETHKKK
```

**Sequence alignment of the four C-terminal Zinc Fingers of Zelda orthologs**

#### Supplementary Figure 7

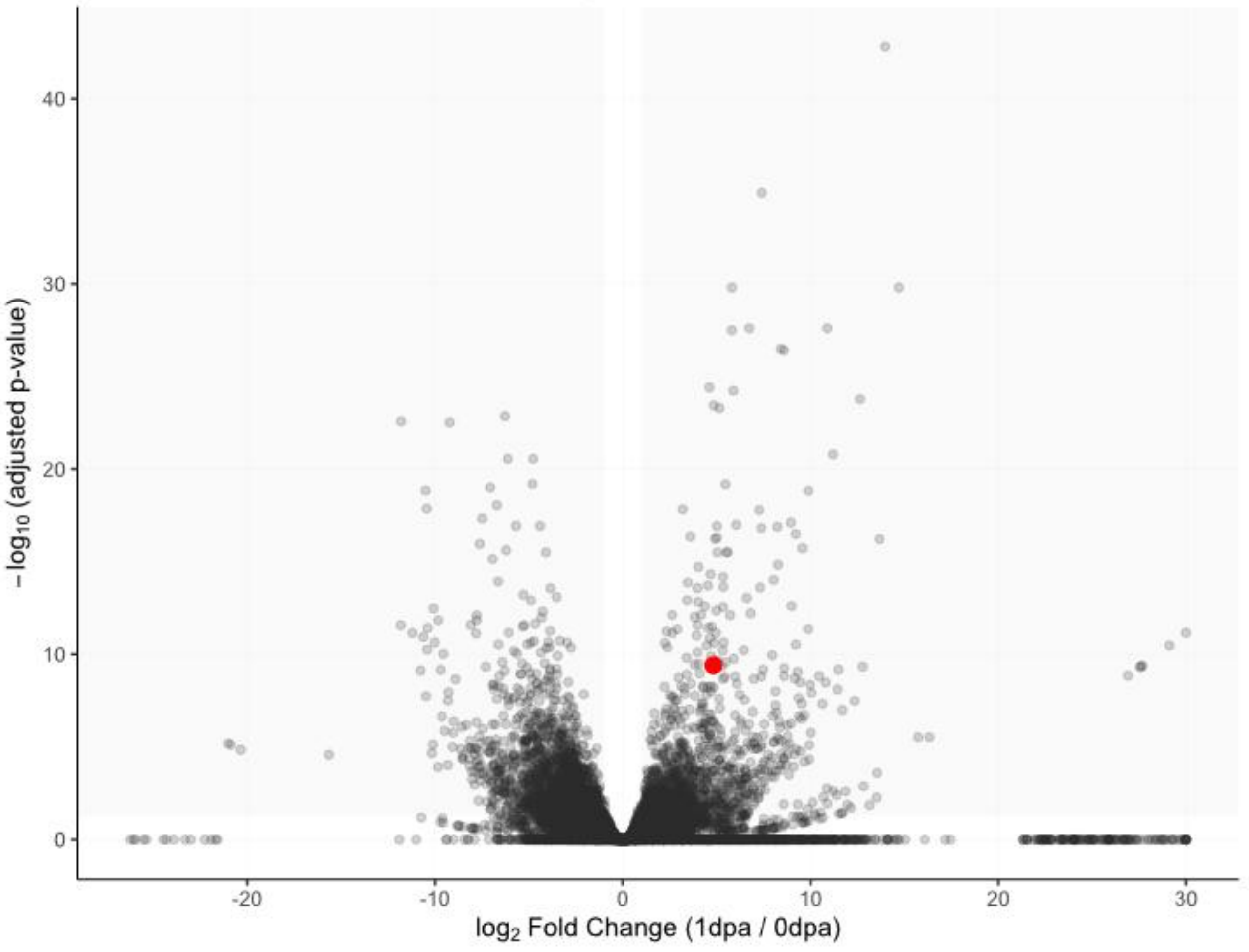


Unzoomed version of Fig. 5F.

#### Supplementary Table 1

| Sample | Total reads | FRiP | Number of peaks | Downsampled total reads | FRiP (downsampled) | Number of peaks (downsampled) |
|---|---|---|---|---|---|---|
| 0dpa_R1 | 20,233,184 | 0.32 | 75,485 | 10,000,000 | 0.28 | 52,832 |
| 0dpa_R2 | 16,508,456 | 0.18 | 48,577 | 10,000,000 | 0.17 | 40,178 |
| 0dpa_R3 | 10,206,744 | 0.21 | 48,936 | 10,000,000 | 0.21 | 47,340 |
| 1dpa_R1 | 39,623,078 | 0.098 | 80,111 | 10,000,000 | 0.044 | 13,573 |
| 1dpa_R2 | 41,257,742 | 0.10 | 85,777 | 10,000,000 | 0.044 | 13,680 |
| 1dpa_R3 | 47,399,164 | 0.13 | 122,403 | 10,000,000 | 0.045 | 14,506 |

Overview of the total number of reads, peaks called, and fraction of reads that fall into those peaks (FRiP) before and after downsampling to 10 million reads.
